# Mitogenome sequences of domestic cats demonstrate lineage expansions and dynamic mutation processes in a mitochondrial minisatellite

**DOI:** 10.1101/2023.06.13.544779

**Authors:** Emily Patterson, Gurdeep Matharu Lall, Rita Neumann, Barbara Ottolini, Chiara Batini, Federico Sacchini, Aiden P. Foster, Jon H. Wetton, Mark A. Jobling

**Affiliations:** Department of Genetics & Genome Biology, University of Leicester, University Road, Leicester LE1 7RH UK; IDEXX Laboratories Italia S.r.l., Via Guglielmo Silva, 36-20149 Milano (MI), Italy; Bristol Veterinary School, University of Bristol, Langford House, Langford, North Somerset, BS40 5DU, UK

**Keywords:** Domestic cat, mitochondrial DNA, haplogroup, control region, tandem repeat, Nanopore sequencing

## Abstract

As a population genetic tool, mitochondrial DNA is commonly divided into the ∼1-kb control region (CR), in which single nucleotide variant (SNV) diversity is relatively high, and the coding region, in which selective constraint is greater and diversity lower, but which provides an informative phylogeny. In some species, the CR contains variable tandemly repeated sequences that are understudied due to heteroplasmy. Domestic cats (*Felis catus*) have a recent origin and therefore traditional CR-based analysis of populations yields only a small number of haplotypes. To increase resolution we used Nanopore sequencing to analyse 119 cat mitogenomes via a long-amplicon approach. This greatly improves discrimination (from 15 to 87 distinct haplotypes) and defines a phylogeny showing similar starlike topologies within all cat haplogroups, likely reflecting post-domestication expansion. We sequenced RS2, a CR tandem array of 80-bp repeat units, placing RS2 array structures within the phylogeny and increasing overall haplotype diversity. Repeat number varies between 3 and 12 (median: 4) with over 30 different repeat unit types differing largely by SNVs. Five SNVs show evidence of independent recurrence within the phylogeny, and seven are involved in at least 11 instances of rapid spread along repeat arrays within haplogroups. In defining mitogenome variation our study provides key information for the forensic genetic analysis of cat hair evidence, and for the first time a phylogenetically informed picture of tandem repeat variation that reveals remarkably dynamic mutation processes at work in the mitochondrion.

## Introduction

Mitochondrial DNA (mtDNA) has long been a widely used tool for the genetic analysis of animal populations [1, 2] because of its unusual properties compared to the nuclear genome: a relatively high mutation rate that contributes to high diversity; a lack of recombination that allows a simple phylogeny to be constructed; and uniparental inheritance that provides information about female population histories. Its high copy number also makes it useful in animal forensic applications [3, 4], with highly degraded or trace materials often yielding mtDNA data when nuclear DNA analysis is not possible.

Most animal mitochondrial genomes are 16-17 kb in size (Figure 1) with ∼95% of this being genic, typically comprising 13 protein-coding, 22 tRNA, and two rRNA genes. However, one segment, the ∼1-kb control region (CR) includes no coding elements, instead containing sequences that control replication and transcription, and the site of the D-loop, a triple-stranded region formed by stable incorporation of a short third DNA strand (7S DNA; [5]). Constraint on sequence evolution in the CR is weaker than in the coding region and consequently it shows higher sequence diversity [6, 7], making it a traditional focus of mtDNA diversity studies. As well as single-nucleotide variation, in many species the CR contains tandemly-repeated sequences [8, 9] that can vary in copy number among individuals. Such sequences provide a potentially useful source of variation, but interpretation is complicated by the fact that they tend to display heteroplasmy [9] - the presence of multiple different length variants within a cell, tissue or individual.

**Figure 1:**
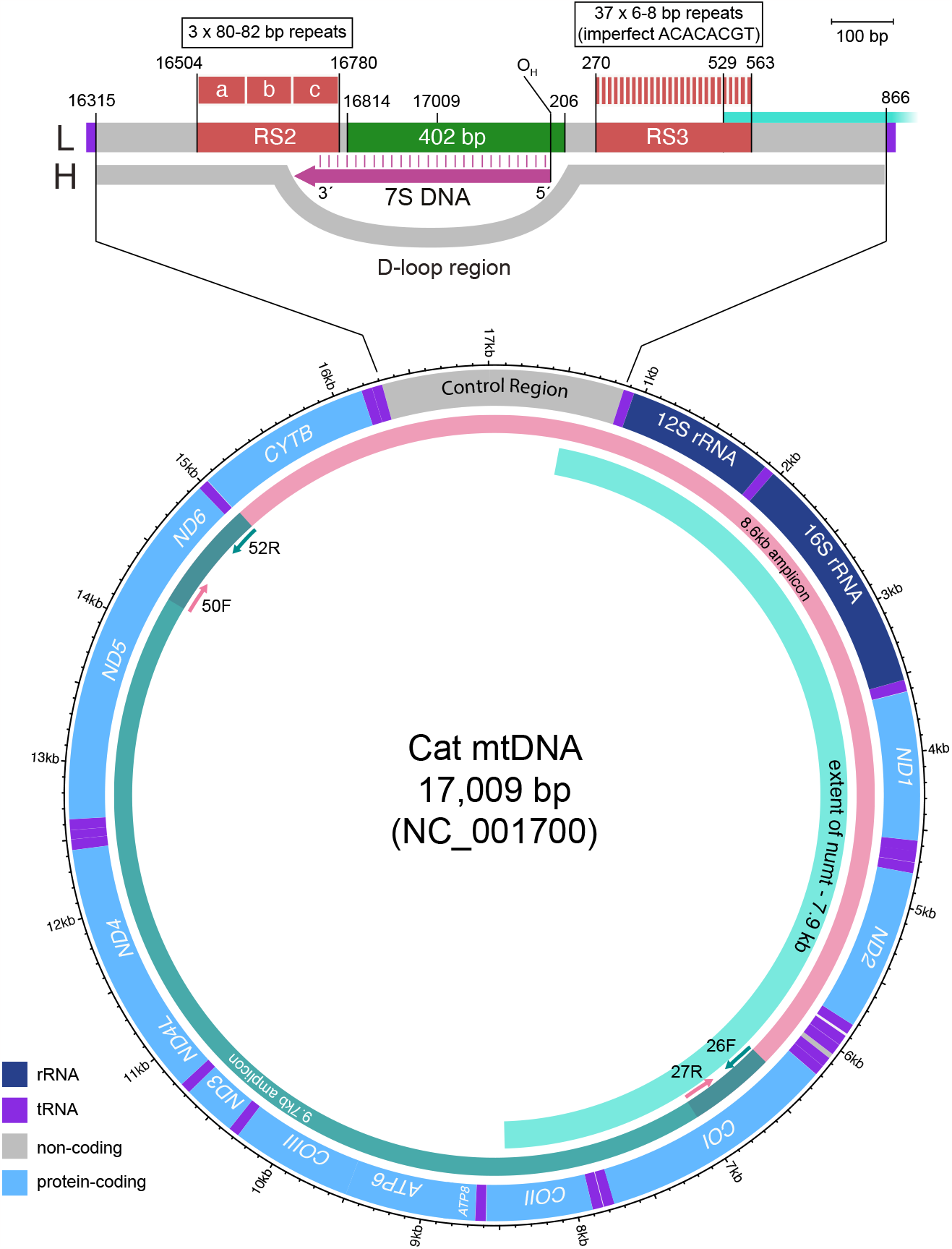
Organisation of the domestic cat mitogenome. Structure of cat mtDNA based on the NC_001700 reference sequence [16], showing genes and the extent of the large numt. Primers for the two long overlapping amplicons used here are indicated by short coloured arrows. Above is an expanded view of the control region, including the positions of the RS2 and RS3 repeat arrays, the 402-bp segment usually sequenced, the origin of heavy strand replication (OH), the D-loop and the 7S DNA. L: light strand; H: heavy strand.

Sequencing the entire molecule is clearly the best approach to maximising the information from mtDNA and over time this approach has been overtaking CR-based studies. One strategy is to design multiple primer pairs that amplify the mitogenome as a set of short overlapping fragments that can be sequenced either using Sanger technology or, more recently, massively parallel sequencing methods [10]. However, a problem with this approach is the presence in all animal nuclear genomes of numts (nuclear mitochondrial segments [11, 12]), some of which exist in high copy number. If primer sequences match numt copies, then these may be amplified alongside, or instead of, the true mtDNA, providing misleading information.

Maximising the detection of mtDNA diversity is particularly important in domesticated animals, whose histories are dominated by founder events and bottlenecks reflecting small numbers of domestication events and selective breeding. The domestic cat (*Felis catus*) has a relatively recent history, with the earliest claimed evidence for domestication dating to 9.5 KYA (thousand years ago) in Cyprus [13], followed by more extensive evidence from Egypt some 4000 years later [14]. Population studies focusing on a 402-bp region of the CR (Figure 1) identify twelve haplogroups (‘mitotypes’) designated A-L, with just four of these (A-D) representing 60-70% of individuals in worldwide cat populations [15]. The sequenced segment is flanked by tandem repeat arrays known as RS2 and RS3 [16]; in the cat mtDNA reference sequence (NC_001700 [16]) RS2 contains three copies of a ∼80-82-bp repeat, while RS3 contains 37 copies of a 6-8-bp repeat. Polymorphism and heteroplasmy have been reported for RS2 in big cats [17] and for RS3 in domestic cats [18]. This complex variability leads to these sequences being generally avoided in diversity studies. An additional complicating factor in domestic cat mitogenome sequencing is the presence on nuclear chromosome D2 of a tandemly repeated 7.9-kb numt (Figure 1) that corresponds to a mitochondrial segment spanning from the CR 3’ end to the *COII* gene [11].

The amplification of mtDNA as two large overlapping amplicons via long PCR provides a means to avoid common multi-copy numts, by the judicious placement of primer sites. The resulting amplicons are also amenable to sequencing using long-read technologies such as Nanopore sequencing, which provides a cost-effective way to survey whole mitogenome variation. In this study, we use this technology to generate whole mitogenome sequences for a population of domestic cats, providing a greater understanding of diversity, illuminating RS2 repeat sequence evolution in a phylogenetic context, and contributing data on the potential discriminatory power of mitogenomes in forensic applications.

## Materials and Methods

### DNA samples

This research was approved by the University of Leicester’s Animal Welfare and Ethical Review Body (ref. AWERB/2021/159) and all blood samples were taken by veterinary surgeons during routine clinical examination. Samples from 119 domestic cats included 115 from a set of 152 individuals from 105 veterinary surgeries throughout England as described previously [19]; the remaining four were from the Bristol Veterinary School. Samples from a wildcat (*Felis silvestris*) and a sand cat (*F. margarita)* were respectively from Twycross Zoo and our laboratory collection. Table S1 lists all studied individuals, their sources and the analyses undertaken. In sub-sampling the 115 cats we used CR-based mitotype information from a previous Sanger sequencing study [19], including less common mitotypes as well as multiple individuals of the same mitotype. A population differentiation test [36] conducted between the full set of 152 and the 119 studied here indicated that the haplotype frequencies did not differ significantly (p=0.98); hence, the dataset presented here is representative. DNA was extracted from 200 μl of blood using the QIAamp DNA Mini Kit (Qiagen), and quantified using the NanoDrop 2000 (Thermo Scientific).

### Long-PCR and sequencing of mtDNA

The mitogenomes of 93 cats (of which 89 were from a previous study [19]), were each amplified in two overlapping segments of 8670 and 9789 bp using primer pairs 26F (5’- AATCGTCACTGCCCATGC-3’) and 52R (5’-TCGAATGTTGGTCATTAAGTT-3’), and 50F (5’- AATGAGCCTAACTATGAGCCA-3’) and 27R (5’-CCATGTAGCCAAAGGGTTC-3’) respectively (Figure 1a). Primer design was based on the domestic cat mitochondrial reference (NC_001700 [16]). Primers were custom barcoded with a 13-bp 5’ sequence and unique barcode combinations were used (Table S2). Barcodes were designed [20] to differ from each other even when accounting for Nanopore sequencing errors.

Amplification was carried out in a 10 μl volume containing 20 ng of template DNA, 0.3 μM of each primer, 0.9 μl of ‘11.1x buffer’ [21] and 0.06 μl of a 20:1 mix of PCRBIO Taq Polymerase (PCRBiosystems; 5U/μl): Pfu DNA polymerase (Agilent Technologies; 2.5 U/μl). PCR cycling conditions for the amplification of cat mitogenome segments generally consisted of an initial denaturation at 96°C for 1 min, followed by 28 cycles of denaturation at 96°C for 20 s, annealing at 60°C for 30 s, extension at 68°C for 9 min for the 8.6-kb segment or 10 min for the 9.7-kb segment, followed by a final extension at 68°C for 10 min.

PCR products were quantified by agarose gel electrophoresis. Equimolar concentrations of the two overlapping amplicons from up to eight cats, each amplified with primers bearing unique custom barcode combinations, were pooled prior to secondary barcoding at the library preparation stage. Each pool was then purified with a 1.0x volume of AppMag PCR Clean Up Beads (Appleton Woods Ltd.) and purified products quantified using the dsDNA HS Assay kit with the Qubit fluorometer 2.0. A Nanopore DNA library was prepared according to the ligation sequencing amplicons - native barcoding amplicon protocol (ONT version: NBA_9093_v109_revF_12Nov2019) using the native barcoding expansion 1-12 (EXP- NBD104) and ligation sequencing kit (SQK-LSK109). Sequencing was carried out using a MinION flow cell (FLO-MIN106, R9.4.1) with a Mk1b MinION sequencer and the Software MinKNOW (v20.10.3) without the basecalling option. Mitogenome fragments from a wildcat and sand cat were amplified (without custom barcodes), quantified and purified as above. A DNA library was prepared following the native barcoding genomic DNA protocol for Flongle (ONT version: NBE_9065_v109_revY_14Aug2019) although the the NEBNext FFPE DNA repair mix and buffer were excluded during the DNA repair and End-prep stage. Sequencing was undertaken using a Flongle flow cell (FLO-FLG001) with the MinKNOW software (v20.10.3).

A total of 35 cat mitogenomes were sequenced using Illumina technology. Amplification used the same PCR reaction mixture and cycling conditions as above, except that 35 cycles were used rather than 28. For 34 samples, sequencing was carried out in-house using the Nextera XT DNA Library Preparation Kit PE 150-bp Illumina MiSeq [MiSeq Reagent Cartridge v2]. One sample (cat67; HgH) was sequenced by Novogene, Cambridge, to a read-depth of 60,337x on a NovaSeq 6000 SP platform. Nine cat mitogenomes were sequenced using both Nanopore and Illumina sequencing technologies to test concordance.

### Processing of sequence data

In processing Nanopore data, Guppy (v.5.0.16) was used for basecalling using the high accuracy model with Qscore filtering disabled and for demultiplexing of Nanopore library native barcodes. Demultiplexing was performed based on the presence of the native barcode on both ends of a read for MinION flow cell runs, and based on the presence of one barcode only for Flongle sequencing data (unless stated otherwise). Read quality statistics were assessed using NanoPlot (v.1.33.1) [22] and reads filtered using NanoFilt (v2.8.0) [22].

Demultiplexing of reads based on our PCR custom barcodes was carried out before filtering. MiniBar (v 0.21) [23] was used to search for barcode and primer sequences within the first and last 150 bp of any given read, with these sequences then being trimmed. Up to two nucleotide differences were allowed in the barcode sequences and up to 11 in the primer sequences. NanoFilt (v2.8.0) [22] was used to remove reads with a quality score of <11, and a length of <6000 or >16,000 bp. Two hundred reads were subsampled for each amplicon for the flow cell sequencing data. Two domestic cats yielded <200 reads for the 9.7-kb amplicon, and therefore all reads were retained.

The two overlapping amplicons were merged and mapped to the domestic cat reference mtDNA sequence (NC_001700) using minimap2 (v2.17) [24] (for the domestic cat, *Felis catus*; NC_028310 for the wildcat, *F. silvestris*; and NC_028308 for the sand cat, *F. margarita*). SAMtools (v1.13) [25] was used to process the mapping files and variants called using freebayes (v0.9.21)[26] with a ploidy of 1. SNPs were then filtered using a quality (QUAL) cut-off of 200. A consensus sequence was generated using BCFtools [27] and the resulting sequences were combined with the Illumina-generated cat mitogenome sequences and aligned using MUSCLE [28]. Illumina sequence data generated on the MiSeq platform were processed as previously described [29] and variants were called using BCFtools mpileup with the -B flag and VarScan (v.2.3.9) mpileup2cns [30].

### Data analysis

Sequence data were visualised using IGV, the Integrative Genomics Viewer [31]. ShinyCircos [32] was used to draw circular representations of mtDNA features and summary data. The effects of variants within coding regions were assessed using the Ensembl Variant Effect Predictor (v105) [33]. Maximum parsimony trees were constructed using PHYLIP v3.698 [34]. Three trees were created using DNAPARS, where the input order of sequences was randomised with a different seed number each time and jumbled ten times. A consensus tree was created using the Consense package and this was used to obtain the final tree in DNAPARS. Median-joining networks [35] were constructed using Network 10 (fluxus-engineering.com) represented using Network Publisher. Population differentiation testing was carried out within Arlequin ver 3.5.2.2 [36]. Dating of clades in the mtDNA phylogeny was done in Network using the mean mutational distance to the relevant root haplotype (rho; [37]). We considered only the coding region and estimated mutation rate based on a mean sand cat/domestic cat divergence time of 1.92 MYA (range: 0.39 - 4.64 MYA [38]), assuming a constant evolutionary rate. The estimated mutation rate was 4.65E-09 substitutions per bp per year (range: 1.92E-09 - 2.29E-08). Secondary structure of RS2 repeats was assessed using mfold [39].

## Results

To assess whole mitogenome variation in domestic cats we undertook Nanopore sequencing of two overlapping mtDNA amplicons (8.6 and 9.7 kb in length) in blood DNA samples from 93 unrelated individuals (Table S1). Altogether, sequencing generated 3.2 million reads, of which ∼60% were successfully allocated to one of 12 native barcodes; of those reads, 43% were then demultiplexed based on their custom barcodes, allowing assignment to individual cats. After filtering for quality and read-length, a total of 706,528 reads were taken for further analysis (an average of 5709 reads per individual for the 8.6-kb amplicon [minimum: 842 reads], and an average of 1887 reads for the 9.7-kb amplicon [minimum: 166]). We also generated 35 mitogenome sequences using the same PCR primers via Illumina sequencing; nine mitogenomes were sequenced with both technologies. A consensus sequence was generated for each mitogenome and the Nanopore sequences were combined with the Illumina-generated sequences and aligned.

Although Nanopore sequencing provides access to long reads, a disadvantage of the LSK109 chemistry used for this project was its relatively high error rate [40], particularly within homopolymeric runs. Among the 93 domestic cats, ten apparent single-base deletions were identified within homopolymeric runs that were 5-8 nucleotides in length according to the reference sequence (Table S3a). Four of these lie within protein-coding regions and, if genuine, would represent frameshift mutations, suggesting that they are artefacts; all four were absent from the Illumina domestic cat sequences. Homopolymer variation in Illumina data suggests that five of the non-coding single-base deletions observed in Nanopore data may be genuine (Table S3a) but to be conservative we exclude all such deletions from analysis. With the exception of the homopolymer indels, and the repetitive RS2 and RS3 regions (discussed below) a general comparison between Nanopore and Illumina data showed complete concordance.

### Single-nucleotide variation within the cat mitogenome assessed by long-amplicon sequencing

In considering sequence variation in the cat mitogenomes, we excluded the repetitive region RS3 (Figure 1a) since its short (6-8-bp) repeat units and overall length (∼300 bp in the reference sequence) make it difficult to interpret (Figure S1). In the reference sequence, the repetitive region RS2 (Figure 1a) contains fewer, longer repeat units (three copies of an 80-82 bp element). However, our sequence data demonstrate that this array contains variation both in repeat sequence and copy number which we consider in the following section. In this section, we restrict our attention to single-nucleotide variation in the mitogenome excluding RS2 and RS3: we can therefore compare variants among cats over a total of 16,438 bp (coordinates: 1-269, 564-16503, 16781-17009).

To understand the variants in our generated sequences we aligned them to the cat reference sequence. In doing this we noted 40 base substitutions and one indel that appear fixed in our sequence dataset compared to the reference. These either represent true private variants or errors within the reference sequence, which was generated by Sanger technology [16]. The reference-specific variants show an unexpectedly strong (17:23) bias towards transversions, and 15/29 coding region variants among them represent apparent missense mutations (Table S4), which supports the idea that they are sequencing artefacts.

The high observed concordance between Nanopore and Illumina mitogenome sequence data allows us to combine the two datasets and consider this as a unified set of 119 cat mitogenomes. Our Nanopore variant calling approach considers the majority nucleotide at any position and therefore is insensitive to heteroplasmy in the sequence data; we therefore do not consider heteroplasmy in this analysis.

The 119 mitogenomes contain a total of 438 variant sites (Table S5), of which 435 are base substitutions and three are indels. The indels (not in homopolymeric runs) are listed in Table S3b, with all three lying in non-coding regions. Among base substitutions, the transition: transversion ratio for the whole mitogenome is 19.45, and that for the control region (coordinates 16315-16503, 16781-269, 564-865) is 4.87. These values are similar to those found in dogs [41], where a ratio of 18.22 was observed in the mitogenome (excluding a VNTR), although a higher transition:transversion ratio of 10.86 was seen in the dog control region [42]. The 333 base substitutions within protein-coding genes include 270 synonymous and 63 missense variants; analysis with VEP indicates that none of these missense mutations is likely to have an impact on protein function that is more than moderate (data not shown). Figure 2 shows the locations of different classes of variants.

**Figure 2:**
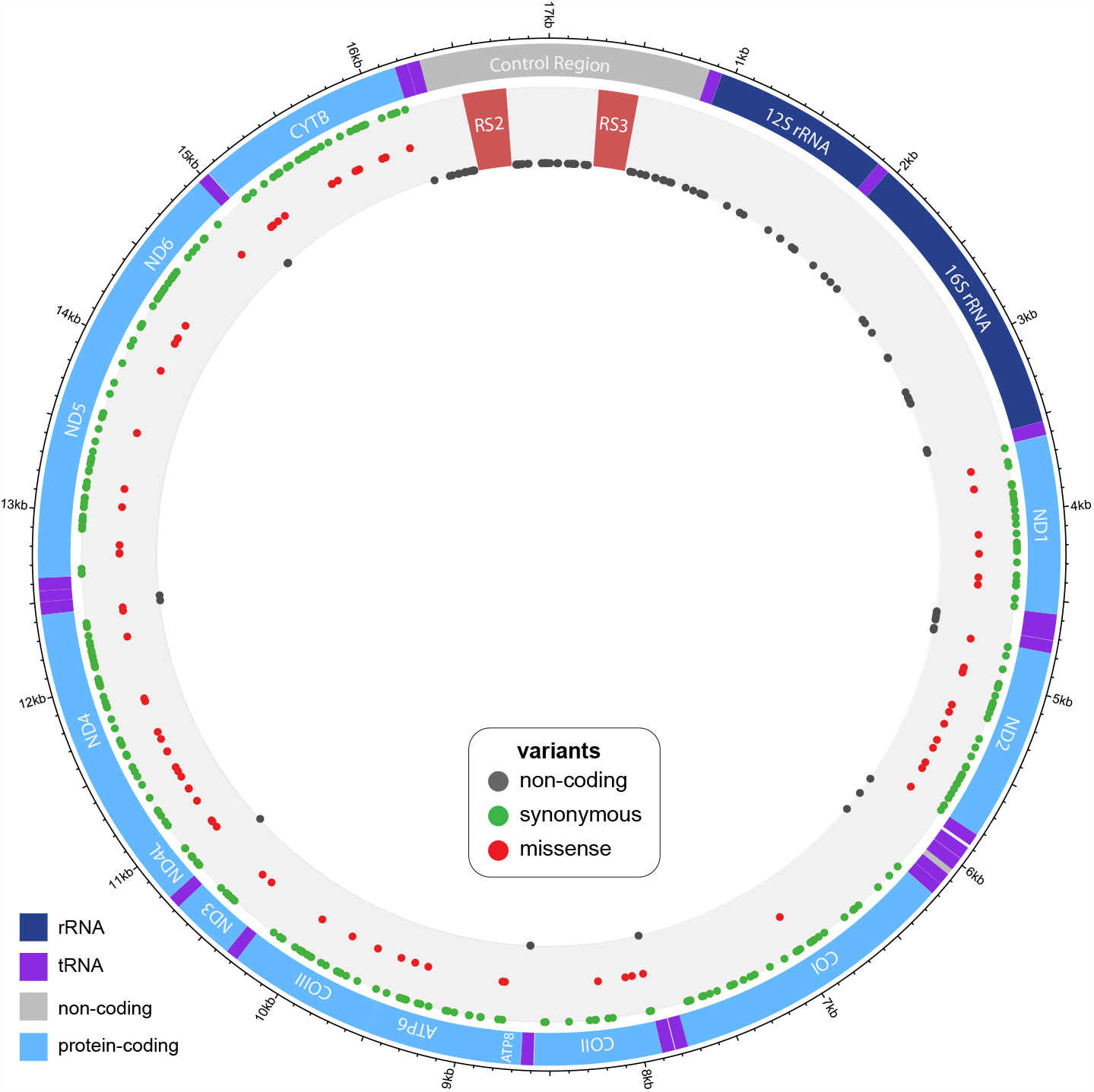
Observed mtDNA sequence variants within the dataset. The 438 variant sites (435 base substitutions and three indels) in mitogenome sequence data from 119 domestic cats, plus the variants in the NC_001700 reference sequence [16] are indicated as filled circles in grey (within non-protein-coding regions), green (synonymous variants in protein-coding genes) or red (missense variants). Variants within the RS2 and RS3 repeat regions are not considered here.

### Cat mitogenome phylogeny

The 119 domestic cat mitogenome sequences, plus the reference sequence NC_001700 and the wildcat sequence generated here, can be represented in a maximum parsimony tree (Figure 3a; Figure S2), rooted to a sand cat (*F. margarita*) outgroup (a median-joining network is shown in Figure S3). The tree also includes the domestic cat reference sequence: as discussed above, there is good reason to believe that this sequence contains errors, which are likely responsible for its lying on a very elongated terminal branch (Figure S2).

**Figure 3:**
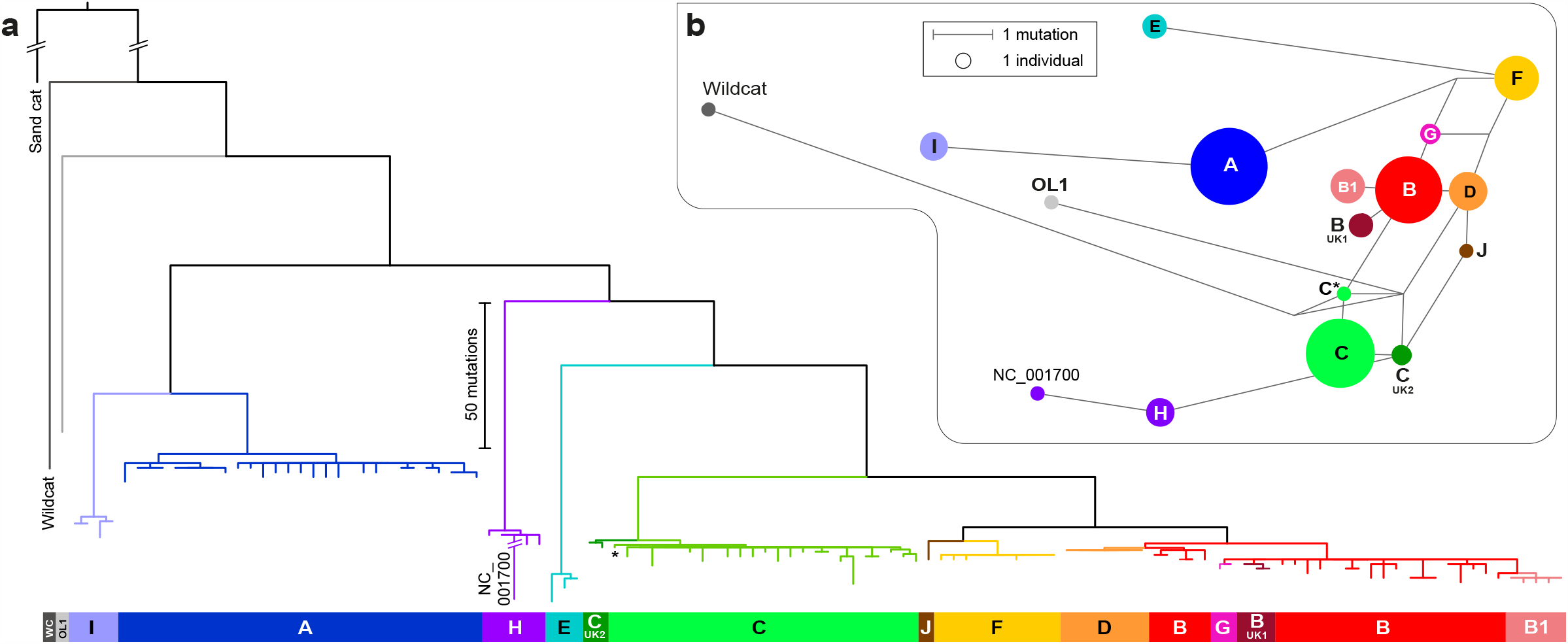
Maximum parsimony phylogeny based on 119 cat mitogenome sequences compared to the CR variant network. a) Maximum parsimony tree rooted to a sand cat mitogenome outgroup (*Felis margarita*; NC_028308) with the addition of a wild cat mitogenome and the *F. catus* reference sequence (NC_001700; [16]). Clades are labelled with the original (A-L) haplogroup names based on control region variant sites. Note the long (here, truncated) terminal branch leading to the NC_001700 reference sequence, which reflects likely sequencing errors. Figure S2 shows the same tree including sample names. b) Median-joining network based on variants within the control region in the 119 cat mitogenomes, plus the wild cat and NC_001700 reference sequence [16]. Circles represent haplotypes, with area proportional to sample size, and lines between haplotypes represent mutational steps as shown in the key.

Clades within the tree are named according to the haplogroups previously defined on the basis of CR sequence variation [15]. The single ‘OL1’ mitogenome, defined as a rare ‘outlier’ haplotype based on CR analysis [15, 19], lies basally in the tree and is more closely related to the wildcat sequence than to domestic cat sequences. This seems consistent with an origin via introgression from a wildcat population. All haplogroup-level clades are monophyletic, and CR-based definitions are highly congruent with the phylogenetic structure of the mitogenome-based tree.

CR-defined haplogroups were based on a relatively small number of variants; the addition of many more variants here shows that, while most established clades (A, C, E, H, I, J and F) are phylogenetically reasonable, some are not. For example, hgs B1, B-UK1 and G are completely embedded within the larger terminal portion of hg B and do not deserve special status based on our tree (Figure 3a), and hg D and the smaller basal portion of hg B form a clear unit.

Among the 119 sequenced mitogenomes there are 87 distinct sequences; haplogroup D is notable in containing seven mitogenomes that show no observable sequence diversity.

While 85% of our sampled cats of known breed are domestic short- or long-hair cats, three of the hg D individuals belong to the Burmese or the related Tonkinese breed, which in the western world are thought to derive from a female imported from Burma in 1930 [43]. This could therefore reflect a recent founder effect; no other breed-haplogroup relationships are evident in the dataset.

The deep structure of the cat mitogenome phylogeny contains no features indicative of ancient expansions (Figure 3a): internal branches are long, involving no polytomies. The tips, however, present a highly consistent appearance showing star-like patterns of similar mutational depths in all lineages, compatible with domestication-related expansions across the entire tree. To estimate the expansion times of these clades we need a measure of mutation rate. The sand cat/domestic cat split is dated at 1.92 MYA (0.39 - 4.64 MYA) [38] which, based on the 276 substitutions in the 15,449-bp coding region between sand cat reference sequence and domestic cat (using cat67, closely related to the reference sequence but without its putative errors) implies a mutation rate of 4.65E-09 (1.93E-09 - 2.29E-08) per base per year. We used the rho statistic [37], i.e. the mean number of mutations to the root, to estimate the average clade expansion date. For the coding region, average rho for the star-like expansion clades A, B/D, C, E, F, H and I is 2.24, which equates to an average clade age of 31.2 KYA (6.3 - 75.3 KYA). We expect the expansions to post-date the onset of domestication, which, given the archaeological evidence, has been placed between 9.5 and 5.7 KYA [13, 14]. This period lies well towards the lower end of our age estimate range, suggesting that our phylogenetic mutation rate estimate may be artificially low; this could reflect issues with the fossil record, hybridisation between lineages [38] or domestication-related relaxation of selection on mtDNA, as suggested for domestic dogs [44].

### Variation within the RS2 repeat array

Within the cat mitochondrial reference sequence [16] the RS2 repeat array contains three copies of an ∼80-bp repeat unit (Figure 4a). Alignment of long-amplicon sequence reads to the reference sequence across this region revealed evidence that many of the sequenced mitogenomes contain different numbers of repeat units (data not shown); this included apparent mixed nucleotides produced through misalignment, similar to the RS3 region (Figure S1).

**Figure 4:**
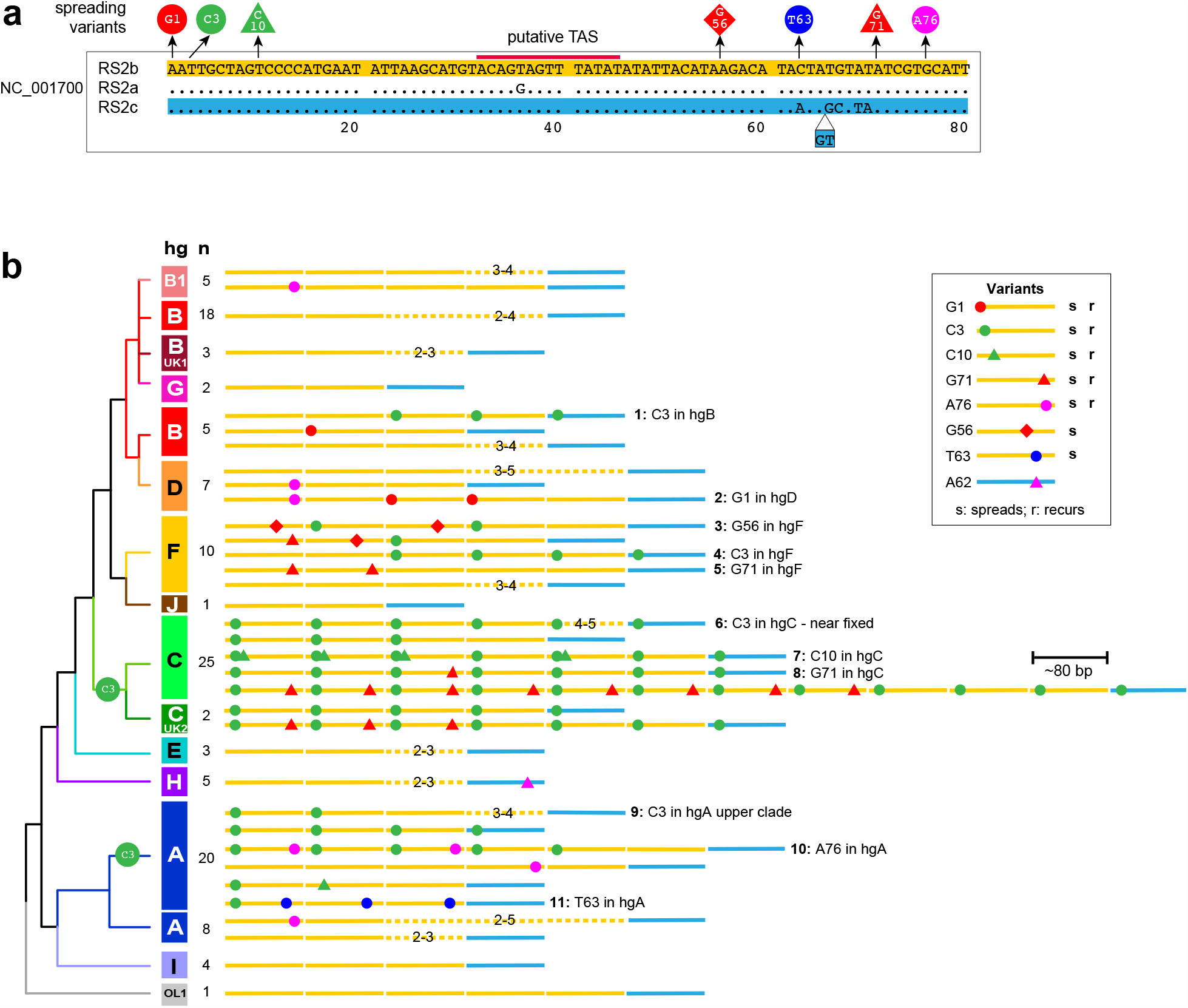
Sequence variation in the RS2 repeat region. a) The three RS2 repeats within the reference sequence (NC_001700). Seven single-nucleotide variants that spread within RS2 arrays are shown, as is the position of a putative termination associated sequence (TAS; [17]). b) Simplified phylogeny showing examples of RS2 repeat array structures and the eleven inferred variant spreading events (labelled 1-11). The full set of 118 analysed arrays can be seen in Figure S5 and Table S6.

To investigate this variation, we amplified a smaller segment of mtDNA encompassing RS2 (755 bp based on the reference sequence) and analysed the products by agarose gel electrophoresis in a subset of cats (Figure S4). Each cat presented heterogeneous PCR fragments of different lengths and intensities, with periodicity around 80 bp; given the length of the repeat unit and the consistency of amplification product lengths between replicate PCRs (data not shown), it seems unlikely that PCR slippage is responsible for the observed heterogeneity and it more probably represents heteroplasmic variation in copy number of the RS2 repeat unit among mitogenome copies. The length of the predominant PCR fragment varied among cats, suggesting underlying repeat unit copy number variation in RS2.

By filtering sequence reads on the basis of binned lengths separated by ∼80 bp, it was possible to capture pools of reads corresponding to PCR products of different lengths and RS2 repeat copy numbers (Figure S4). This allowed the repeat copy number of the major molecular species in each cat to be identified and the sequence determined using a reference-free approach (Supplementary methods). In this way we could analyse both copy number and sequence of RS2 repeat regions in 118/119 of the Nanopore-sequenced cat mitogenomes and place this repeat array variation in the context of the phylogeny (Figures 4 and S5; Table S6).

Within the cat reference mtDNA sequence, the first two repeats (80 bp; designated RS2a and b [16]) differ by one base substitution, while the third (82 bp; RS2c) is somewhat divergent, containing five base substitutions and a 2-bp insertion in its last 20 bp (Figure 4a). The 118 sequenced RS2 arrays follow the same general pattern (Figure 4b, S5; Table S6), with a single RS2c-like repeat preceded by a set of between 2 and 11 repeat unit copies of RS2b-like repeats; overall median repeat number is 4. The RS2b repeat is found in all haplogroups of the phylogeny including the deepest rooting haplotype, OL1, and seems likely to represent the ancestral repeat type. However, among RS2b-like repeats across the phylogeny there are 30 variant repeat sequences that differ largely by single-nucleotide variants. Nine such variants are singletons, but the remainder are shared between haplotypes in a way that shows clear evidence of multiple independent recurrence of variants, and the dynamic diffusion of variants along repeat arrays within haplogroups.

Examples of such variant spreading events are highlighted in Figure 4b, and the full set of RS2 haplotypes is illustrated in Figure S5 and described in Table S6. A maximum parsimony approach indicates that there have been at least 11 different variant spreading events within the phylogeny.

Additional haplotype discrimination obtained by adding unambiguous RS2 variants contributed by SNVs in the 118 cats to the sequence variation in the rest of the molecule (Table S5) increases the number of haplotypes from 87 different haplotypes based on SNV variation (Figures 3; S2) to 103, and this is further increased to 110 haplotypes when repeat array length is considered.

## Discussion

Cats are widely believed to have become domesticated in the Near East through association with the development of agriculture (since 12 KYA) and the concomitant increase in rodent pests linked to farming and food storage. Dates from archaeological studies are consistent with this, the earliest claimed evidence dating to 9.5 KYA [13]. Subsequent human migration and trade are likely to have spread domesticated cats widely and rapidly, and this is reflected in a lack of feline population structure as detected by analysis of multiple short-tandem repeat and single-nucleotide polymorphisms in a large sample of mostly Eurasian cats [45]. Likewise, mtDNA CR haplotypes are widely shared among populations, albeit at different frequencies [19] and a few common mitotypes predominate.

Here we have used Nanopore and Illumina sequencing of long PCR amplicons to analyse the mitogenomes of 119 domestic cats, producing a highly resolved SNP-based phylogeny based on 16,438 bp of sequence. The deep structure of the phylogeny (Figure 3a) contains no polytomies and no evidence of ancient expansions of domestic cat matrilineages. One basal haplotype (OL1) represents a possible wildcat introgression. The phylogeny contains five haplogroups (A, C, E, H and I) that are separated by deep splits; haplogroups in the remaining well-separated superhaplogroup (encompassing B, D, G, F and J) are more closely related. Within clades, we observe star-like topologies of similar depths, which suggests domestication-related expansions across the tree. However, dating based on the median sand cat/domestic cat divergence time produces unexpectedly ancient expansion ages, suggesting possible issues with the fossil dates, discordance due to ancient hybridisation (widespread among the Felidae [38]) or recent increases in effective mutation rates via relaxed selection [44]. One notable feature in the phylogeny is the extended branch leading to the reference sequence (NC_001700); we interpret this as evidence of errors in the original sequence, and suggest use of an amended reference as an alternative, based on the cat67 mitogenome, which is the most closely related sequence to NC_001700 in our phylogeny (Figure S2).

Vertebrate mitochondrial DNA is often portrayed as a paragon of economy, with its conserved and minimal gene set, and its lack of wasteful introns or intergenic sequences. In this context, the presence in some species of repeat arrays at five different positions in the CR [9] is surprising. Diversity in such repeat regions has been understudied because of the problems of heteroplasmy, and past Sanger-based analysis has often relied on cloning [9] which in turn may have introduced artefacts. Current sequencing approaches such as Nanopore technology offer the opportunity to use bioinformatic filtering to examine individual populations of reads [46], and thereby throw light on complex structural variation. The cat RS3 array clearly contains variation in repeat sequence that appears to be phylogenetically structured (Figure S1) as well as variation in copy number, but remains challenging to analyse; here, we have focused our attention on RS2.

Although RS2 variation has previously been studied between related species (e.g. big cats [17] and ruminants [47]) or within a sample of a species (e.g. Japanese sika deer [48]), our study of the structures of the cat RS2 repeat array presents the first (to our knowledge) picture of the diversity of a mitochondrial minisatellite-like sequence within the framework of a detailed intraspecific SNV-based phylogeny. This allows events that are identical by descent (ancestral) or identical by state (recurrent) to be distinguished, and provides evidence of a remarkably dynamic process of recurrent mutation and variant spread across arrays. The blood DNA samples studied here show evidence of a high degree of RS2 heteroplasmy in the form of multiple different length variants in individual cats, as observed in PCR products and length profiles of sequence reads (Figure S4). We have chosen to focus on the apparent majority length class in each cat in defining a representative RS2 array structure. However, this is not entirely objective and majority length could vary between tissues.

What can be deduced about underlying RS2 mutation processes? In early descriptions of mitochondrial repeat diversity (focused on RS3; [9]), repeat secondary structures in single-stranded DNA were invoked as drivers of intrahelical misalignment and replication slippage. This is also implied in the original description of cat RS2 [16], which illustrated a secondary structure involving two repeats. However, the structures presented were for the folding of RNA, rather than DNA; in terms of free energy, RNA secondary structures are much more stable than those of equivalent DNA sequences [49, 50] because of the additional hydrogen bonds permitted by ‘wobble’ base pairing and by the presence of the 2’-hydroxyl group of ribose. Reanalysis of RS2 using DNA-folding [39], by contrast, produces structures that show very weak intramolecular interactions (Figure S6) which seem unconvincing candidates for secondary structure formation. The RS2 repeat unit contains [16, 17] a putative termination associated sequence (TAS), which has been linked to the termination of replication of the 7S DNA molecule (Figure 1; [5]) and in human cells is reversibly bound by a helicase, TWINKLE [51]. The TAS was first identified in human-mouse comparisons [52] and lies some distance rightwards (in our orientation; Figure 1) of the 3’ end of the 7S DNA, as assessed from direct analyses of this molecule [52, 53]. It therefore seems likely that much of the heavy strand of the control region, including some RS2 repeat units, is single-stranded while the D-loop structure exists. If this were important in the generation and spread of new repeat variants in the RS2 array, it might be expected to lend some polarity to repeat array variation; however, the structures shown in Figures 4 and S5 do not display this.

The long-amplicon based mitogenome sequencing undertaken here greatly increases observed sequence diversity compared to the standard approach of CR analysis. The 16,438 bp sequenced (excluding RS2 and RS3) in 119 cats yields 87 distinct sequences, a marked increase from the 15 observed CR haplotypes. This represents a reduction of random match probability (RMP) from 0.16 to 0.018, and therefore it illustrates that mitogenome sequencing will have forensic utility. Since most forensic casework involves cat hairs [19, 54], in which DNA is likely to be degraded and long-amplicon approaches unsuccessful, it will be necessary to develop suitable short-amplicon multiplexes (as described elsewhere; Patterson et al., in prep.). If it were possible to add RS2 sequence diversity to assessments of forensic evidence, this would further decrease RMP, as has been suggested for RS3 [18].

However, the high degree of observed heteroplasmy is likely to present problems in the rigorous definition of matches or exclusions.

## Data availability

Sequences described in this paper have been submitted to GenBank and accession numbers are listed in Table S1. Sequences are also provided as fasta files and an alignment at https://figshare.com/articles/dataset/Domestic_cat_mtDNA_sequences/23266712.

## Acknowledgments

EP was supported by a BBSRC-MIBTP (grant no. BB/M01116X/1) iCASE studentship, co-sponsored by Twycross Zoo (East Midland Zoological Society) and Zoological Society of London. We thank Jane and Andrew Elliot for additional financial support. We gratefully acknowledge colleagues who contributed DNA samples (including Angie Hibbert and colleagues from the Langford Feline Centre and Jade Ward from Bristol Veterinary School) and NUCLEUS Genomic Services at the University of Leicester for access to Illumina MiSeq sequencing. This research used the SPECTRE High Performance Computing Facility at the University of Leicester for data analysis.

## References

[1] R.G. Harrison, Animal mitochondrial DNA as a genetic marker in population and evolutionary biology, Trends Ecol Evol 4 (1989) 6–11.

[2] M.W. Bruford, D.G. Bradley, G. Luikart, DNA markers reveal the complexity of livestock domestication, Nat Rev Genet 4 (2003) 900–10.

[3] S. Kanthaswamy, Review: domestic animal forensic genetics - biological evidence, genetic markers, analytical approaches and challenges, Animal Genet 46 (2015) 473–84.

[4] A. Linacre, Animal Forensic Genetics, Genes (Basel) 12 (2021).

[5] T.J. Nicholls, M. Minczuk, In D-loop: 40 years of mitochondrial 7S DNA, Exp Gerontol 56 (2014) 175–81.

[6] G.G. Brown, G. Gadaleta, G. Pepe, C. Saccone, E. Sbisa, Structural conservation and variation in the D-loop-containing region of vertebrate mitochondrial DNA, J Mol Biol 192 (1986) 503–11.

[7] C. Saccone, G. Pesole, E. Sbisa, The main regulatory region of mammalian mitochondrial DNA: structure-function model and evolutionary pattern, J Mol Evol 33 (1991) 83–91.

[8] L. Fumagalli, P. Taberlet, L. Favre, J. Hausser, Origin and evolution of homologous repeated sequences in the mitochondrial DNA control region of shrews, Mol Biol Evol 13 (1996) 31–46.

[9] A.R. Hoelzel, J.V. Lopez, G.A. Dover, S.J. O’Brien, Rapid evolution of a heteroplasmic repetitive sequence in the mitochondrial DNA control region of carnivores, J Mol Evol 39 (1994) 191–9.

[10] W. Parson, G. Huber, L. Moreno, M.B. Madel, M.D. Brandhagen, S. Nagl, C. Xavier, M. Eduardoff, T.C. Callaghan, J.A. Irwin, Massively parallel sequencing of complete mitochondrial genomes from hair shaft samples, Forensic Sci Int Genet 15 (2015) 8–15.

[11] J.V. Lopez, N. Yuhki, R. Masuda, W. Modi, S.J. O’Brien, Numt, a recent transfer and tandem amplification of mitochondrial DNA to the nuclear genome of the domestic cat, J Mol Evol 39 (1994) 174–90.

[12] F.M. Calabrese, D.L. Balacco, R. Preste, M.A. Diroma, R. Forino, M. Ventura, M. Attimonelli, NumtS colonization in mammalian genomes, Sci Rep 7 (2017) 16357.

[13] J.D. Vigne, J. Guilaine, K. Debue, L. Haye, P. Gerard, Early taming of the cat in Cyprus, Science 304 (2004) 259.

[14] W. Van Neer, V. Linseele, R. Friedman, B. De Cupere, More evidence for cat taming at the Predynastic elite cemetery of Hierakonpolis (Upper Egypt), J Archaeol Sci 45 (2014) 103–111.

[15] R.A. Grahn, J.D. Kurushima, N.C. Billings, J.C. Grahn, J.L. Halverson, E. Hammer, C.K. Ho, T.J. Kun, J.K. Levy, M.J. Lipinski, J.M. Mwenda, H. Ozpinar, R.K. Schuster, S.J. Shoorijeh, C.R. Tarditi, N.E. Waly, E.J. Wictum, L.A. Lyons, Feline non-repetitive mitochondrial DNA control region database for forensic evidence, Forensic Sci Int Genet 5 (2011) 33–42.

[16] J.V. Lopez, S. Cevario, S.J. O’Brien, Complete nucleotide sequences of the domestic cat (Felis catus) mitochondrial genome and a transposed mtDNA tandem repeat (Numt) in the nuclear genome, Genomics 33 (1996) 229–46.

[17] K. Jae-Heup, E. Eizirik, S.J. O’Brien, W.E. Johnson, Structure and patterns of sequence variation in the mitochondrial DNA control region of the great cats, Mitochondrion 1 (2001) 279–92.

[18] F. Fridez, S. Rochat, R. Coquoz, Individual identification of cats and dogs using mitochondrial DNA tandem repeats?, Sci Justice 39 (1999) 167–71.

[19] B. Ottolini, G.M. Lall, F. Sacchini, M.A. Jobling, J.H. Wetton, Application of a mitochondrial DNA control region frequency database for UK domestic cats, Forensic Sci Int Genet 27 (2017) 149–155.

[20] A. Srivathsan, E. Hartop, J. Puniamoorthy, W.T. Lee, S.N. Kutty, O. Kurina, R. Meier, Rapid, large-scale species discovery in hyperdiverse taxa using 1D MinION sequencing, BMC Biol 17 (2019) 96.

[21] A.J. Jeffreys, R. Neumann, V. Wilson, Repeat unit sequence variation in minisatellites: a novel source of DNA polymorphism for studying variation and mutation by single molecule analysis, Cell 60 (1990) 473–485.

[22] W. De Coster, S. D’Hert, D.T. Schultz, M. Cruts, C. Van Broeckhoven, NanoPack: visualizing and processing long-read sequencing data, Bioinformatics 34 (2018) 2666–2669.

[23] H. Krehenwinkel, A. Pomerantz, J.B. Henderson, S.R. Kennedy, J.Y. Lim, V. Swamy, J.D. Shoobridge, N. Graham, N.H. Patel, R.G. Gillespie, S. Prost, Nanopore sequencing of long ribosomal DNA amplicons enables portable and simple biodiversity assessments with high phylogenetic resolution across broad taxonomic scale, Gigascience 8 (2019).

[24] H. Li, Minimap2: pairwise alignment for nucleotide sequences, Bioinformatics 34 (2018) 3094–3100.

[25] H. Li, B. Handsaker, A. Wysoker, T. Fennell, J. Ruan, N. Homer, G. Marth, G. Abecasis, R. Durbin, The Sequence Alignment/Map format and SAMtools, Bioinformatics 25 (2009) 2078–2079.

[26] E. Garrison, G. Marth, Haplotype-based variant detection from short-read sequencing, arXiv 1207.3907 (2012).

[27] P. Danecek, A. Auton, G. Abecasis, C.A. Albers, E. Banks, M.A. DePristo, R.E. Handsaker, G. Lunter, G.T. Marth, S.T. Sherry, G. McVean, R. Durbin, The variant call format and VCFtools, Bioinformatics 27 (2011) 2156–2158.

[28] R.C. Edgar, MUSCLE: multiple sequence alignment with high accuracy and high throughput, Nucleic Acids Res 32 (2004) 1792–7.

[29] C. Batini, P. Hallast, A.J. Vagene, D. Zadik, H.A. Eriksen, H. Pamjav, A. Sajantila, J.H. Wetton, M.A. Jobling, Population resequencing of European mitochondrial genomes highlights sex-bias in Bronze Age demographic expansions, Sci Rep 7 (2017) 12086.

[30] D.C. Koboldt, K. Chen, T. Wylie, D.E. Larson, M.D. McLellan, E.R. Mardis, G.M. Weinstock, R.K. Wilson, L. Ding, VarScan: variant detection in massively parallel sequencing of individual and pooled samples, Bioinformatics 25 (2009) 2283–5.

[31] J.T. Robinson, H. Thorvaldsdottir, W. Winckler, M. Guttman, E.S. Lander, G. Getz, J.P. Mesirov, Integrative genomics viewer, Nat Biotechnol 29 (2011) 24–26.

[32] Y. Yu, Y. Ouyang, W. Yao, shinyCircos: an R/Shiny application for interactive creation of Circos plot, Bioinformatics 34 (2018) 1229–1231.

[33] W. McLaren, L. Gil, S.E. Hunt, H.S. Riat, G.R. Ritchie, A. Thormann, P. Flicek, F. Cunningham, The Ensembl Variant Effect Predictor, Genome Biol 17 (2016) 122.

[34] J. Felsenstein, PHYLIP (Phylogeny Inference Package) version 3.6. Distributed by the author (Department of Genome Sciences, University of Washington, Seattle, WA). 2005.

[35] H.-J. Bandelt, P. Forster, A. Röhl, Median-joining networks for inferring intraspecific phylogenies, Mol Biol Evol 16 (1999) 37–48.

[36] L. Excoffier, H.E. Lischer, Arlequin suite ver 3.5: a new series of programs to perform population genetics analyses under Linux and Windows, Mol Ecol Resour 10 (2010) 564–7.

[37] J. Saillard, P. Forster, N. Lynnerup, H.-J. Bandelt, S. Nørby, mtDNA variation among Greenland Eskimos: the edge of the Beringian expansion, Am J Hum Genet 67 (2000) 718–726.

[38] G. Li, B.W. Davis, E. Eizirik, W.J. Murphy, Phylogenomic evidence for ancient hybridization in the genomes of living cats (Felidae), Genome Res 26 (2016) 1–11.

[39] M. Zuker, Mfold web server for nucleic acid folding and hybridization prediction, Nucleic Acids Res 31 (2003) 3406–15.

[40] C. Delahaye, J. Nicolas, Sequencing DNA with Nanopores: Troubles and biases, PLoS One 16 (2021) e0257521.

[41] S. Verscheure, T. Backeljau, S. Desmyter, Dog mitochondrial genome sequencing to enhance dog mtDNA discrimination power in forensic casework, Forensic Sci Int Genet 12 (2014) 60–8.

[42] A. Duleba, K. Skonieczna, W. Bogdanowicz, B. Malyarchuk, T. Grzybowski, Complete mitochondrial genome database and standardized classification system for Canis lupus familiaris, Forensic Sci Int Genet 19 (2015) 123–129.

[43] Cat Fanciers’ Association, The Tonkinese; https://cfa.org/tonkinese/. 2022).

[44] S. Bjornerfeldt, M.T. Webster, C. Vila, Relaxation of selective constraint on dog mitochondrial DNA following domestication, Genome Res 16 (2006) 990–4.

[45] S.M. Nilson, B. Gandolfi, R.A. Grahn, J.D. Kurushima, M.J. Lipinski, E. Randi, N.E. Waly, C. Driscoll, H. Murua Escobar, R.K. Schuster, S. Maruyama, N. Labarthe, B.B. Chomel, S.K. Ghosh, H. Ozpinar, H.C. Rah, J. Millan, F. Mendes-de-Almeida, J.K. Levy, E. Heitz, M.A. Scherk, P.C. Alves, J.E. Decker, L.A. Lyons, Genetics of randomly bred cats support the cradle of cat domestication being in the Near East, Heredity (Edinb) 129 (2022) 346–355.

[46] C. Marshall, W. Parson, Interpreting NUMTs in forensic genetics: Seeing the forest for the trees, Forensic Sci Int Genet 53 (2021) 102497.

[47] A. Hassanin, A. Ropiquet, A. Couloux, C. Cruaud, Evolution of the mitochondrial genome in mammals living at high altitude: new insights from a study of the tribe Caprini (Bovidae, Antilopinae), J Mol Evol 68 (2009) 293–310.

[48] H. Ba, L. Wu, Z. Liu, C. Li, An examination of the origin and evolution of additional tandem repeats in the mitochondrial DNA control region of Japanese sika deer (Cervus Nippon), Mitochondrial DNA A DNA Mapp Seq Anal 27 (2016) 276–81.

[49] M. Bercy, U. Bockelmann, Hairpins under tension: RNA versus DNA, Nucleic Acids Res 43 (2015) 9928–36.

[50] J.B. Swadling, K. Ishii, T. Tahara, A. Kitao, Origins of biological function in DNA and RNA hairpin loop motifs from replica exchange molecular dynamics simulation, Phys Chem Chem Phys 20 (2018) 2990–3001.

[51] E. Jemt, O. Persson, Y. Shi, M. Mehmedovic, J.P. Uhler, M. Davila Lopez, C. Freyer, C.M. Gustafsson, T. Samuelsson, M. Falkenberg, Regulation of DNA replication at the end of the mitochondrial D-loop involves the helicase TWINKLE and a conserved sequence element, Nucleic Acids Res 43 (2015) 9262–75.

[52] J.N. Doda, C.T. Wright, D.A. Clayton, Elongation of displacement-loop strands in human and mouse mitochondrial DNA is arrested near specific template sequences, Proc Natl Acad Sci U S A 78 (1981) 6116–20.

[53] D.R. Foran, J.E. Hixson, W.M. Brown, Comparisons of ape and human sequences that regulate mitochondrial DNA transcription and D-loop DNA synthesis, Nucleic Acids Res 16 (1988) 5841–61.

[54] L.A. Lyons, R.A. Grahn, T.J. Kun, L.R. Netzel, E.E. Wictum, J.L. Halverson, Acceptance of domestic cat mitochondrial DNA in a criminal proceeding, Forensic Sci Int Genet 13 (2014) 61–7.

