## Supplemental Text for "Mitogenome sequences of domestic cats demonstrate lineage expansions and dynamic mutation processes in a mitochondrial minisatellite"

### **Supplementary Methods**: **Investigation of RS2 region sequence variation**

In order to obtain long-read sequences spanning the entirety of the RS elements, one of three different barcoded versions (CTGAGGTGATCAG, ATCCGGTCGGAGA or AGTGTCCTGCTAG) of primers 57F (CCCATCTCAGGCATTATTG) and 3R (TTTAAGCTACATTAACTGGGA), consisting of the same 13-bp 5´ sequence in both the forward and reverse primer, were used to amplify coordinates 16136-881 of the mitogenomes of the 26 cats which had only been sequenced using Illumina technology. Amplification was performed in a 10 µl volume, containing 10 ng of template DNA, 0.3 µM of each primer, 0.9 µl of “11.1x Buffer” [1], and 0.06 µl of a 20:1 mix of PCRBIO Taq Polymerase (PCRBiosystems; 5 U/µl): Pfu DNA polymerase (Agilent Technologies; 2.5 U/µl). PCR cycling conditions consisted of an initial denaturation at 96°C for 1 min, followed by 28 cycles of denaturation at 96°C for 20 s, annealing at 56°C for 30 s, extension at 68°C for 2 min, and a final extension at 68°C for 25 min. PCR products were quantified by agarose gel electrophoresis, and equimolar concentrations of products from cats with different combinations of forward and reverse barcodes, were pooled together. The twelve pools were purified with a 1.8x volume of AppMag PCR Clean Up Beads (Appleton Woods Ltd.) following the manufacturer’s instructions, and purified products quantified using the dsDNA HS Assay kit with the Qubit 2.0 fluorometer.

After DNA library preparation and sequencing (see main text), Guppy (v.5.0.16) was used for basecalling with Q-score filtering disabled. Demultiplexing of native barcodes was also carried out using Guppy, filtering for the presence of barcodes on both ends of each read. MiniBar (v 0.21) [2] was used for custom barcode demultiplexing based on the presence of the custom barcode within the first and last 150 bp of a given read, with these sequences then trimmed from that read. Up to 11 differences in the primer sequences and two differences in the barcode sequences were allowed. Reads with a quality score of <11 and shorter than 1000 and longer than 3000 bp were removed using NanoFilt (v2.8.0) [3]. The resulting reads were mapped to the domestic cat reference mtDNA sequence (NC_001700) using minimap2 (v2.17) [4], with the mapping files converted to BAM files using SAMtools (v1.13) [5].

At this stage, the filtered, un-subsampled ~9-kb reads from the 93 long-range (Nanopore-sequenced) cats and the 26 cats sequenced above had been aligned to the reference genome. Using the BAM files, SAMtools ampliconclip was used to clip reads using the options `--hard-clip --both-ends --filter-len 0`. Sequences spanning beyond coordinates 16136 -16955 bp for the Flongle-sequenced cats, and coordinates 15591-16955 bp for the MinION-sequenced cats, were clipped. The BAM files were converted back to FASTQ files using BEDTools (v2.28.0) [6]. Read lengths of the FASTQ files were binned by length separated by ~80 bp. The major length species of each cat was used as input with Amplicon_sorter [7] which generated a consensus sequence. If more than one consensus sequence was generated per cat sample, the sequence with the greatest number of supporting reads was retained.

In addition, for a small subset of cats a segment stretching across the RS2 region was amplified and the PCR products visualised by agarose gel electrophoresis. PCR amplifications were carried out in a 10 µl volume, which contained 10 ng of template DNA, 0.3 µM of each of the 58F (CATATATTGCATATACCCATACTG) and 60R (GCTGAGTCATAGCAAGTTCG) primers, 0.9 µl of 11.1x Buffer [1], and 0.06 µl of a 20:1 mix of PCRBIO Taq Polymerase (PCRBiosystems; 5U/µl): Pfu DNA polymerase (Agilent Technologies; 2.5 U/µl). Amplification conditions consisted of an initial denaturation at 96°C for 1 min, followed by 28 cycles at 96°C for 20 s, 62°C for 30 s, 68°C for 1 min, and a final extension at 68°C for 25 min.

**References**

[1] A.J. Jeffreys, R. Neumann, V. Wilson, Repeat unit sequence variation in minisatellites: a novel source of DNA polymorphism for studying variation and mutation by single molecule analysis, Cell 60 (1990) 473-485.
[2] H. Krehenwinkel, A. Pomerantz, J.B. Henderson, S.R. Kennedy, J.Y. Lim, V. Swamy, J.D. Shoobridge, N. Graham, N.H. Patel, R.G. Gillespie, S. Prost, Nanopore sequencing of long ribosomal DNA amplicons enables portable and simple biodiversity assessments with high phylogenetic resolution across broad taxonomic scale, Gigascience 8 (2019).
[3] W. De Coster, S. D'Hert, D.T. Schultz, M. Cruts, C. Van Broeckhoven, NanoPack: visualizing and processing long-read sequencing data, Bioinformatics 34 (2018) 2666-2669.
[4] H. Li, Minimap2: pairwise alignment for nucleotide sequences, Bioinformatics 34 (2018) 3094-3100.

[5] H. Li, B. Handsaker, A. Wysoker, T. Fennell, J. Ruan, N. Homer, G. Marth, G. Abecasis, R. Durbin, The Sequence Alignment/Map format and SAMtools, Bioinformatics 25 (2009) 2078-2079.
[6] A.R. Quinlan, I.M. Hall, BEDTools: a flexible suite of utilities for comparing genomic features, Bioinformatics 26 (2010) 841-2.
[7] A.R. Vierstraete, B.P. Braeckman, Amplicon_sorter: A tool for reference-free amplicon sorting based on sequence similarity and for building consensus sequences, Ecol Evol 12 (2022) e8603.
