## Supplemental Figures for "Mitogenome sequences of domestic cats demonstrate lineage expansions and dynamic mutation processes in a mitochondrial minisatellite"

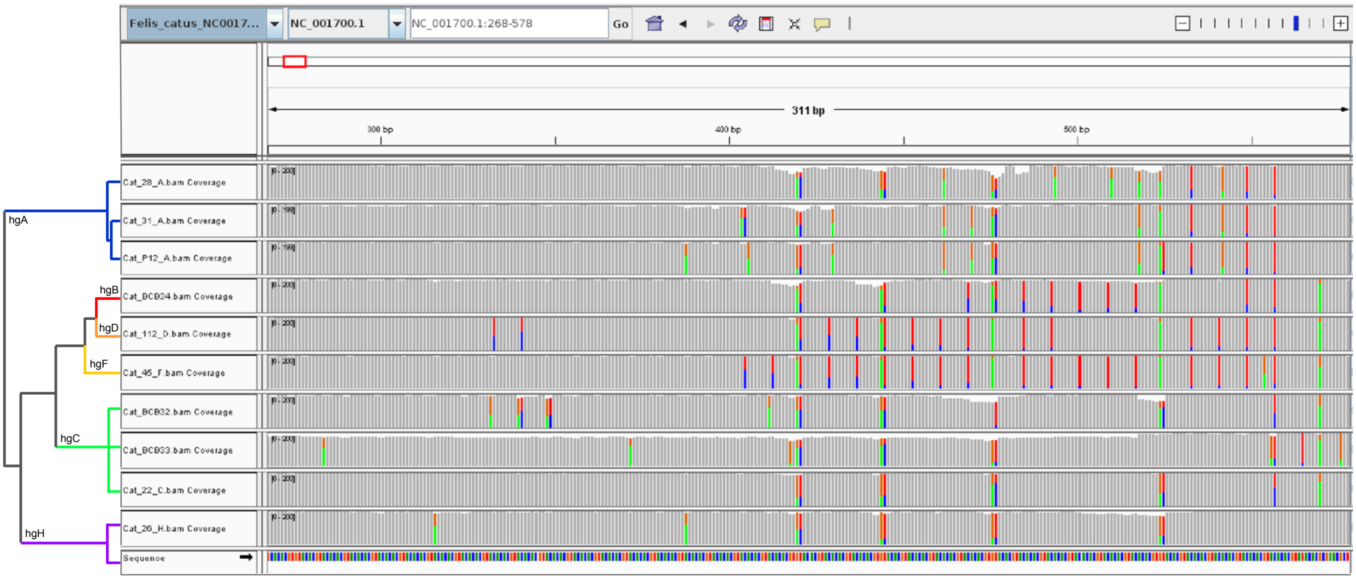


**Figure S1: Complex variation within the RS3 repeat array.**

IGV screenshot showing segments of long-amplicon mtDNA reads from ten randomly selected cats aligned to the domestic cat reference sequence (shown at the bottom of the window). Variants within the RS3 array are visible as differences from the reference (vertical coloured bars) with a periodicity of 6-8 bp. Many variant sites are present as mixed nucleotides (vertical bars of mixed colours), indicating complex differences in repeat copy number and sequence from the reference, and possibly heteroplasmy, resulting in sequence misalignment. This prohibits robust interpretation of variation within RS3. The schematic tree to the left shows relationships between the ten cat mitogenomes, which demonstrates that patterns of variants show phylogenetic coherence.

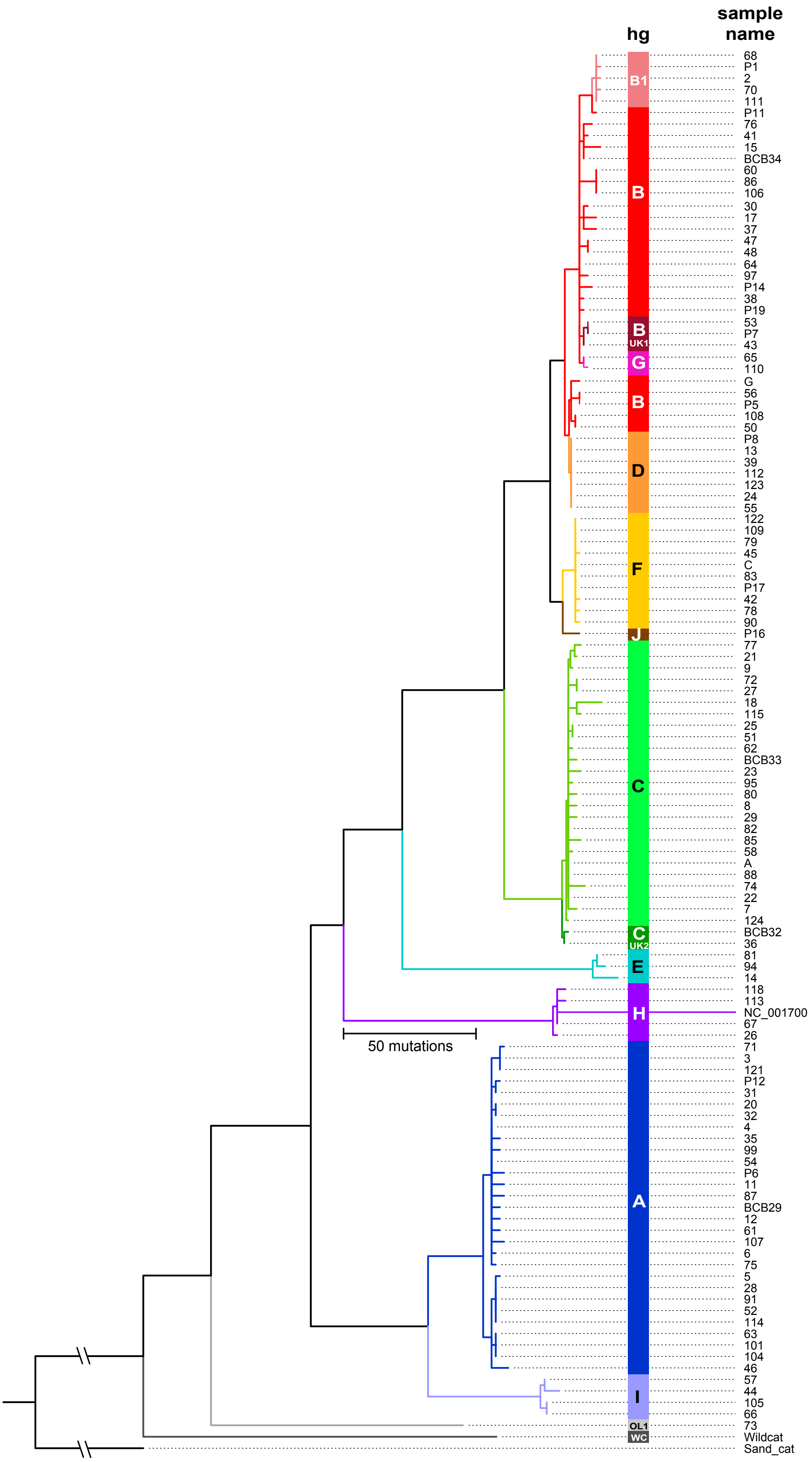


**Figure S2: Maximum parsimony tree based on all mitogenome single-nucleotide variants, including sample names.**

Note the long branch to NC_001700, the cat reference sequence.

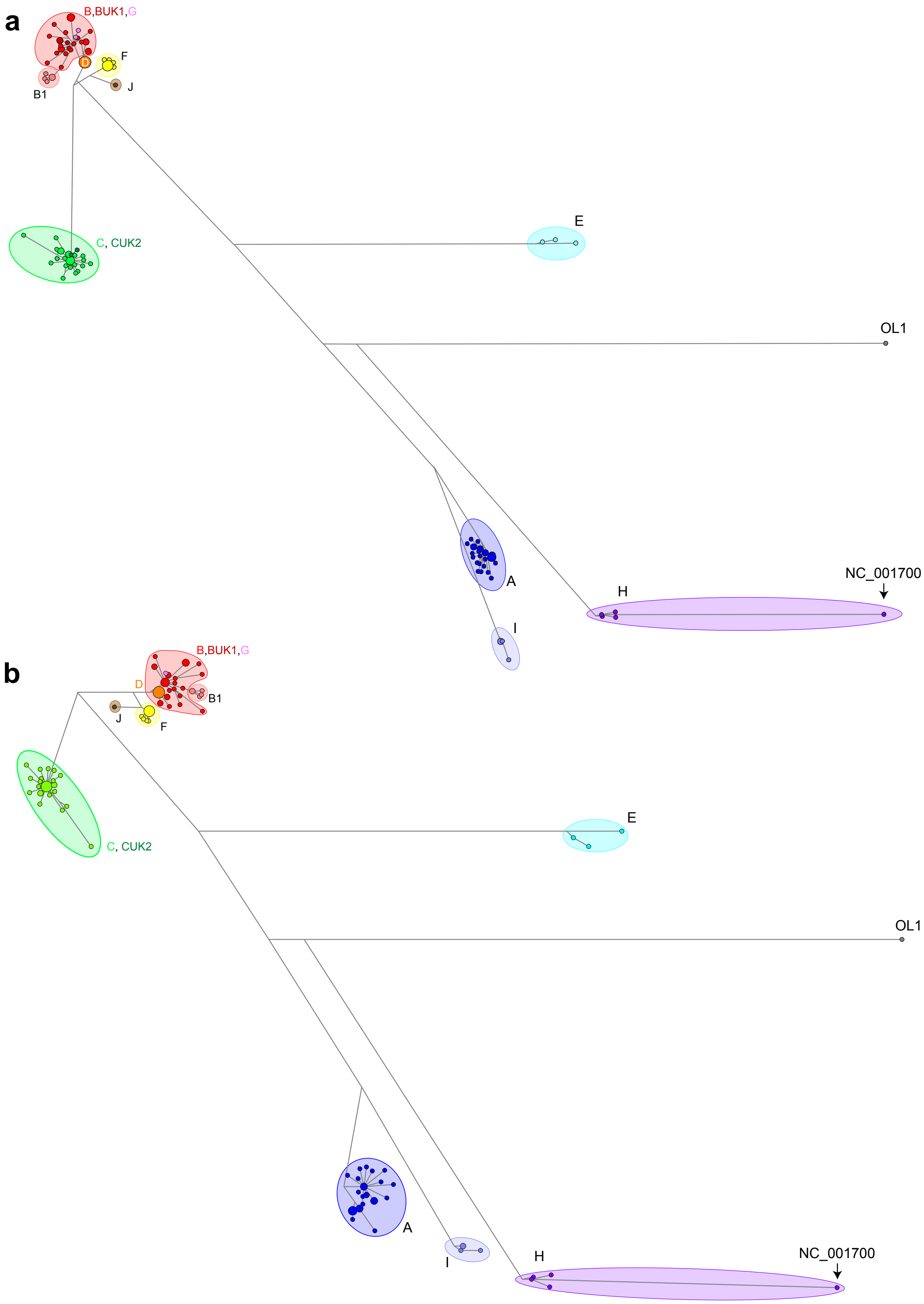


**Figure S3: Median joining networks based on cat mitogenome sequences.**

a) Network based on coding plus control region (16,438 bp; coordinates: 1-269, 564-16503, 16781-17009).
b) Network based on coding region only (15,449 bp; coordinates: 866-16781), as used in rho calculations for dating. Circles indicate haplotypes with area proportional to the number of individuals. Lines between circles represent mutational steps. Haplogroups are indicated by colour shading and labels.

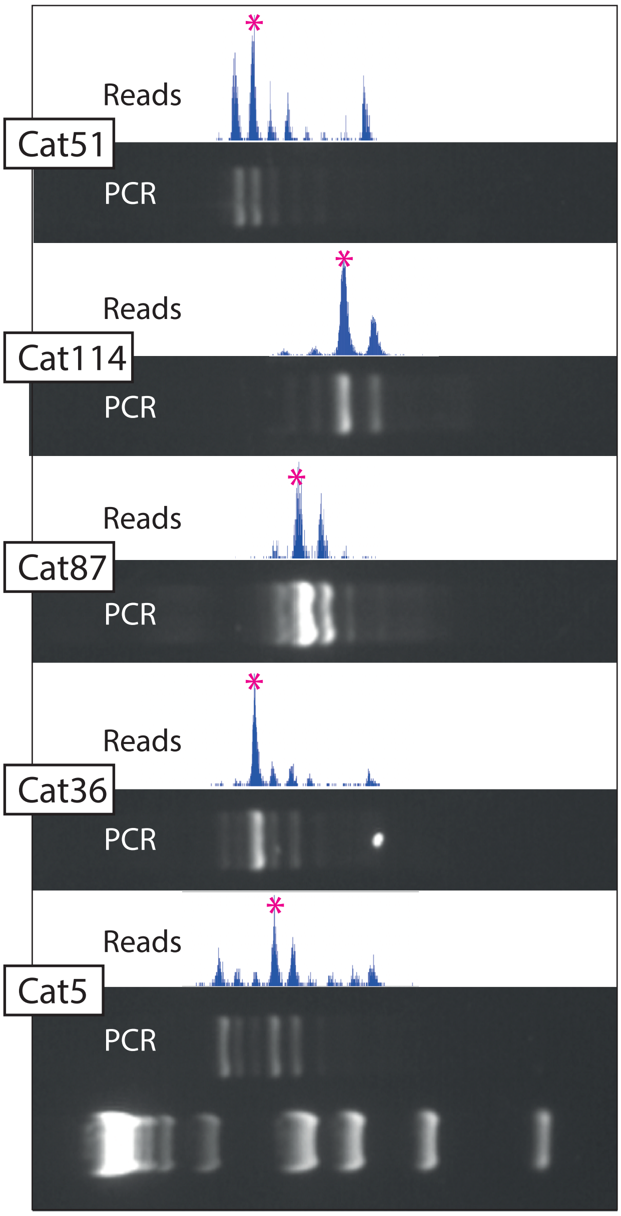


**Figure S4: Correspondence of PCR products and sequence read-depth across the RS2 repeat region in a selection of cats.**

Above each agarose gel image showing PCR products generated across the RS2 repeat region is a plot of sequence read-depth observed in bins separated by ~82 bp. The major bin is highlighted in each case by an asterisk.

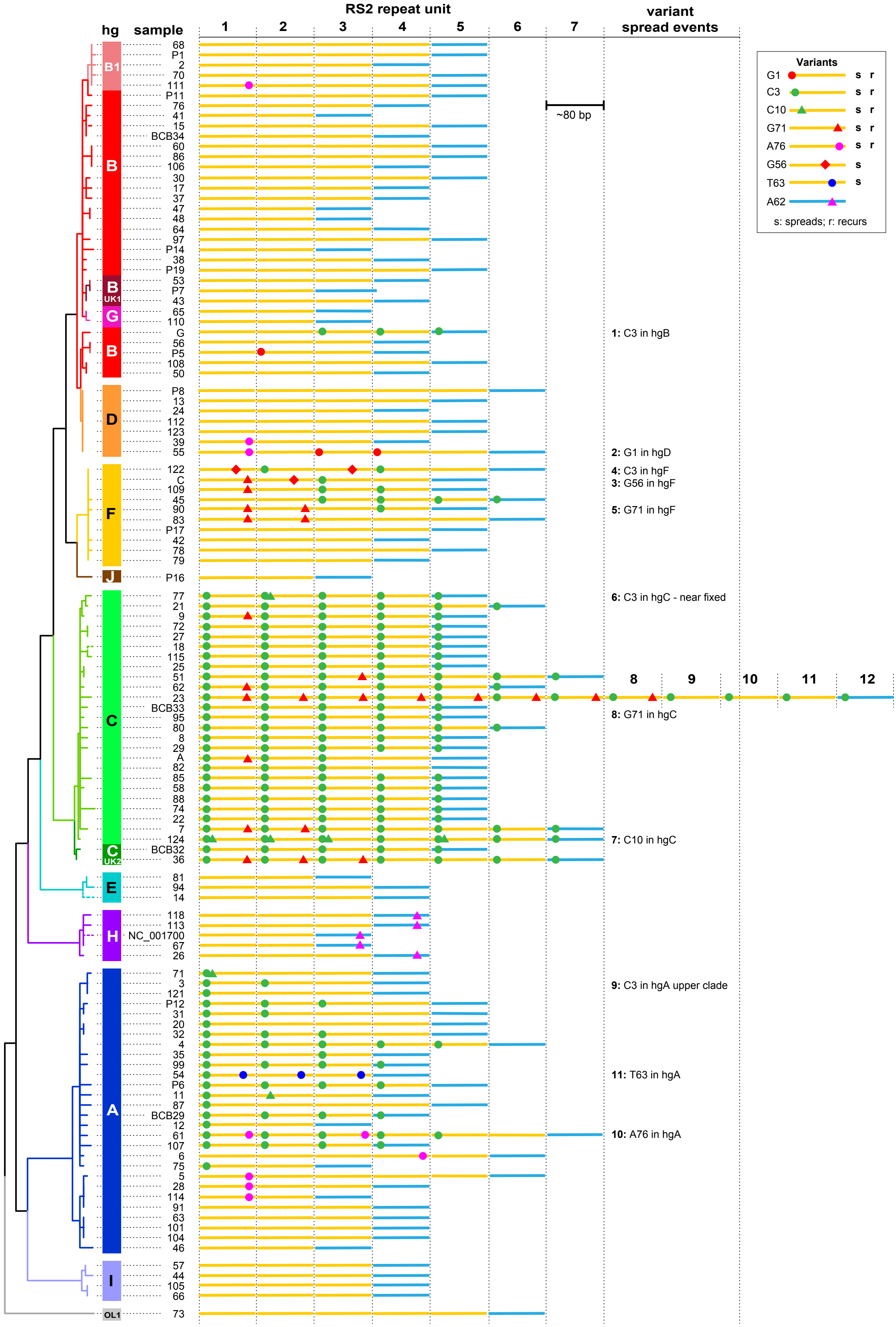


**Figure S5: RS2 repeat variation across domestic cat phylogeny.**

Variants shown are those that recur and/or spread in the repeat array, or singletons that are prevalent or fixed in a haplogroup. Full sequences including singletons can be found in Table S5. Branch lengths in the SNP-based tree to the left are not proportional to mutational distance. Some branches have been moved (compared to Figures 3 and S2) to bring together likely events identical by descent; cat 52 could not be analysed. Spread events are numbered as in Figure 4.

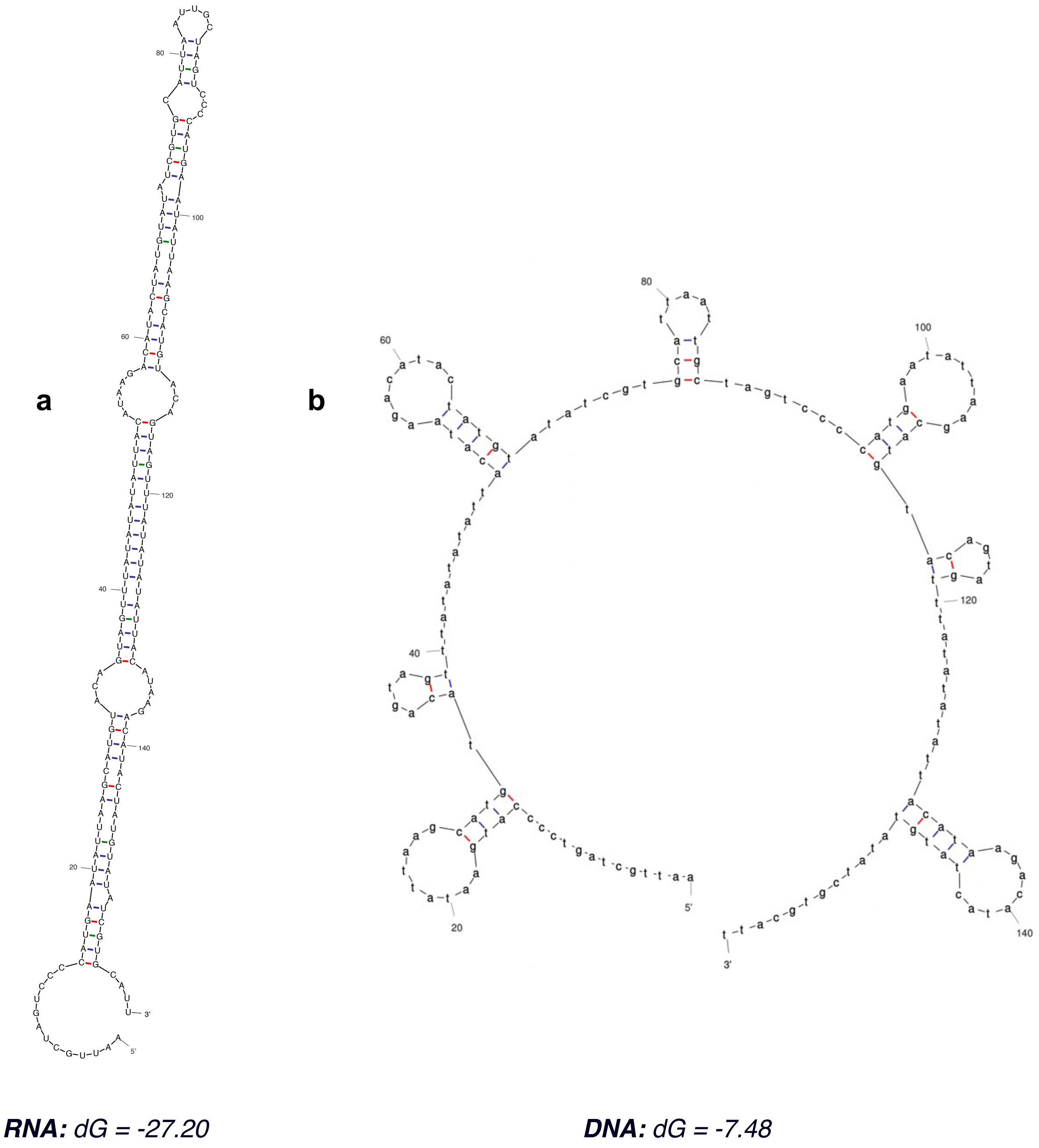


**Figure S6: Folding of RS2 repeats based on RNA and DNA sequences.**

a) Folding of two RS2b repeat types based on RNA, as presented by Lopez et al. (1996).
b) Folding of the same sequence based on DNA.
Analysis was done using mfold (Zuker, 2003). Free energy (dG) of each structure is shown in kcal/mol.
